# Genome-wide screen identifies host loci that modulate *M. tuberculosis* fitness in immunodivergent mice

**DOI:** 10.1101/2023.03.05.528534

**Authors:** Rachel K. Meade, Jarukit E. Long, Adrian Jinich, Kyu Y. Rhee, David G. Ashbrook, Robert W. Williams, Christopher M. Sassetti, Clare M. Smith

## Abstract

Genetic differences among mammalian hosts and *Mycobacterium tuberculosis* (*Mtb*) strains determine diverse tuberculosis (TB) patient outcomes. The advent of recombinant inbred mouse panels and next-generation transposon mutagenesis and sequencing approaches has enabled dissection of complex host- pathogen interactions. To identify host and pathogen genetic determinants of *Mtb* pathogenesis, we infected members of the BXD family of mouse strains with a comprehensive library of *Mtb* transposon mutants (TnSeq). Members of the BXD family segregate for *Mtb*-resistant C57BL/6J (B6 or *B*) and *Mtb*-susceptible DBA/2J (D2 or *D*) haplotypes. The survival of each bacterial mutant was quantified within each BXD host, and we identified those bacterial genes that were differentially required for *Mtb* fitness across BXD genotypes. Mutants that varied in survival among the host family of strains were leveraged as reporters for “endophenotypes”, each bacterial fitness profile directly probing specific components of the infection microenvironment. We conducted QTL mapping of these bacterial fitness endophenotypes and identified 140 *<u>h</u>ost-<u>p</u>athogen* quantitative trait loci (*hp*QTL). We identified a QTL hotspot on chromosome 6 (75.97–88.58 Mb) associated with the genetic requirement of multiple *Mtb* genes; *Rv0127* (*mak*), *Rv0359* (*rip2*), *Rv0955* (*perM*), and *Rv3849* (*espR*). Together, this screen reinforces the utility of bacterial mutant libraries as precise reporters of the host immunological microenvironment during infection and highlights specific host-pathogen genetic interactions for further investigation. To enable downstream follow-up for both bacterial and mammalian genetic research communities, all bacterial fitness profiles have been deposited into GeneNetwork.org and added into the comprehensive collection of TnSeq libraries in MtbTnDB.

## Introduction

Every pathogenic infection is part of an evolutionary battle between host and invader. In the case of *Mycobacterium tuberculosis* (*Mtb*), the causative agent of tuberculosis (TB), evidence tracing back at least 9,000 years tells the tale of a host-pathogen arms race (Hershkovitz et al. 2008), making *Mtb* one of the most enduring adversaries of the human species. Given the prevailing nature of this challenge, it is no surprise that weaknesses in vital host defenses provide greater opportunity for *Mtb* infection, allow *Mtb* to manipulate the nature and magnitude of the host immune response, and worsen patient prognoses (Cooper et al. 1993; Flynn et al. 1993; Cooper et al. 1997; Caruso et al. 1999; Kramnik et al. 2000). In addition, the advent of drug resistant and multi-drug resistant *Mtb* strains highlights the imminent danger posed by our microscopic rivals and underscores the importance of bacterial variation as a critical factor in the pathogenesis and continued spread of *Mtb* (Caws et al. 2008; Hernández-Pando et al. 2012; Gopal et al. 2014). Meanwhile, genetic diversity of both hosts and *Mtb* strains that interact during infection can give rise to a spectrum of disease outcomes (Smith and Sassetti 2018), enhancing the complexity of TB identification and treatment. To combat a pathogen responsible for nearly 10.6 million infections and 1.6 million deaths annually (WHO 2022), it is therefore vital to dissect the host-pathogen interface.

Mammalian models of TB have been used successfully to identify and mechanistically interrogate TB susceptibility loci identified from human cohorts. For mouse strains that are phenotypically divergent, quantitative trait locus (QTL) mapping studies within genetically tractable and reproducible hosts have enabled the identification of genomic loci that contribute to the magnitude of clinically relevant phenotypes (Lavebratt et al. 1999; Kramnik et al. 2000; Mitsos et al. 2000; Mitsos et al. 2003; Yan et al. 2006). The classic inbred strains C57BL/6J (B6) and DBA/2J (D2) lie on opposite ends of the TB susceptibility spectrum, with B6 surviving nearly a year post-infection and D2 succumbing to cachexia and morbidity within months (Medina and North 1998). To capitalize on this diverse disease outcome and just over 6 million divergent SNPs (Wang et al. 2016; Ashbrook et al. 2021; Ashbrook et al. 2022; Sasani et al. 2022), a recombinant inbred panel generated from these two founders, known as the BXD, has been used to understand host determinants of susceptibility to a variety of diseases (Taylor et al. 1973; Taylor et al. 1999; Peirce et al. 2004; Ashbrook et al. 2021).

BXD and other recombinant inbred panels have been used to explore the intricate dynamics of TB disease, but often the phenotypes leveraged to understand these dynamics in mice, such as survival time post-infection, body weight, or bacterial burden, are too complex and polygenic to reveal novel biology or enable refined mapping. These “macrophenotypes” materialize from many genetic and environmental factors that can be difficult to mechanistically dissect or precisely map. Within each unique host, bacteria face similarly diverse immunological dynamics and spatiotemporal selective pressures. To contribute to the growing body of work on genetic determinants of TB outcome, quantifying precise bacterial “endophenotypes” underlying disease outcomes is the crucial next step toward elaborating the biological intricacies that govern the host-pathogen interface. Recently employed molecular endophenotypes include expression QTL (Chesler et al. 2005), metabolite QTL (Wu et al. 2014), protein QTL (Chick et al. 2016), and chromatin accessibility QTL (Skelly et al. 2020), but for infectious disease, the most salient endophenotypes lie at the interface of the host and pathogen.

A classic approach to assess the requirement of a bacterial gene for infection is to infect a standard inbred mouse strain, such as B6, with a single knockout *Mtb* mutant. Expanding on this design, genome-wide mutagenesis enables the generation of single knockout mutants across the *Mtb* genome, thereby allowing the measurement of *Mtb* gene requirements under a variety of selective pressures. Transposon mutant libraries (TnSeq) have been successfully used to report on variable components of the bacterial infection microenvironment under distinct host and antibiotic pressures (Sassetti and Rubin 2003; Nambi et al. 2015; Mishra et al. 2017; Bellerose et al. 2019). We have recently leveraged TnSeq across genetically diverse Collaborative Cross (CC) mice to map both host- and pathogen-linked loci (Smith et al. 2022).

To further characterize the BXD phenome (Chesler et al. 2004) and add to the growing repertoire of selective pressures applied to *Mtb* whole-genome mutant libraries (Jinich et al. 2021), we now leverage the phenotypic divergence and reproducibility of the BXD family and the molecular phenotyping precision of an *Mtb* transposon mutant library. We infected parental genotypes and groups of mice from 19 BXD strains with a saturated library of *Mtb* transposon mutants and quantified mutant abundance within each host at one-month post infection. We conducted dual-genome QTL mapping to identify linkages between bacterial fitness profiles and host genetic factors, revealing 140 host genetic loci significantly associated with specific bacterial mutants. Thus, in passing a bacterial transposon mutant library through selection within the immunologically and phenotypically diverse BXD microenvironments, we have conducted a multidimensional host-pathogen screen to identify precise host and bacterial determinants of tuberculosis disease. Overall, this study provides a blueprint for the functional dissection of host-pathogen interactions that underlie diverse disease outcomes.

## Materials & Methods

### Ethics Statement

All mouse studies were conducted in accordance with the guidelines issued in the Guide for the Care and Use of Laboratory Animals of the National Institutes of Health and the Office of Laboratory Animal Welfare. Animal studies conducted at Duke University Medical School were conducted using protocols approved by the Duke Institutional Animal Care and Use Committee (IACUC) (Animal Welfare Assurance #A221-20-11) in a manner designed to minimize pain and suffering in *Mtb*-infected animals. Any animal exhibiting signs of severe disease was immediately euthanized in accordance with IACUC approved endpoints. All mouse studies conducted at the University of Massachusetts Medical School (UMass) were performed using protocols approved by UMass IACUC (Animal Welfare Assurance #A3306-01).

### Mice

Male and female C57BL/6J (#000664) and DBA/2J (#000671) mice were purchased from The Jackson Laboratory. Male mice from 19 BXD strains were imported from the colony of Robert Williams (University of Tennessee Health Science Center, Memphis, TN) in 2013. The 19 BXD strains in this study include: BXD9/TyJ, BXD29/Ty, BXD39/TyJ, BXD40/ TyJ, BXD48a/RwwJ, BXD51/RwwJ, BXD54/RwwJ, BXD56/RwwJ, BXD60/RwwJ, BXD62/RwwJ, BXD67/RwwJ, BXD69/RwwJ, BXD73/RwwJ, BXD73b/RwwJ, BXD77/RwwJ, BXD79/RwwJ, BXD90/RwwJ, BXD93/RwwJ, BXD102/RwwJ. All mice were housed in a specific pathogen-free facility within standardized living conditions (12-hour light/dark, food and water *ad libitum*). Mice were matched at 8-12 weeks of age at the time of *Mtb* infection. Male mice were used in the TnSeq screen, and both male and female mice were used in the aerosol studies.

### M. tuberculosis *Strains*

All *Mtb* strains were cultured in Middlebrook 7H9 medium supplemented with oleic acid-albumin-dextrose catalase (OADC), 0.2% glycerol, and 0.05% Tween 80 to log-phase with shaking (200 rpm) at 37°C. Hygromycin (50 µg/mL) or kanamycin (20 µg/mL) were added when necessary. Prior to all *in vivo* infections, cultures were washed, resuspended in phosphate-buffered saline (PBS) containing 0.05% Tween 80 (hereafter PBS-T), and sonicated before diluting to desired concentration.

### Mouse Infections

For aerosol infections, B6 and D2 were infected with ∼50 colony forming units (CFU) of *Mtb* H37Rv YFP via aerosol inhalation (Glas-Col) for 6 and 12 weeks. At each timepoint, male and female mice were euthanized in accordance with approved IACUC protocols, and lung and spleen were harvested into PBS-T and homogenized. CFU was quantified by dilution plating onto Middlebrook 7H10 agar supplemented with OADC, 0.2% glycerol, 50 mg/mL Carbenicillin, 10 mg/mL Amphotericin B, 25 mg/mL Polymixin B, and 20 mg/mL Trimethoprim. One lung lobe per mouse was collected into 10% neutral-buffered formalin for H&E staining by the Duke University Research Immunohistology Laboratory.

For TnSeq experiments, 1x10^6^ CFU of saturated *Himar1* transposon mutants (Sassetti et al. 2003) was delivered via intravenous tail vein injection. Mice were infected in a single batch. At 4 weeks post-infection, mice were euthanized, and spleens and lungs were harvested then homogenized in a FastPrep-24 (MP Biomedicals). CFU was quantified by dilution plating on 7H10 agar with 20 µg/mL Kanamycin. For library recovery, approximately 1x10^6^ CFU per mouse was plated on 7H10 agar with 20 µg/mL Kanamycin. After three weeks of growth, colonies were harvested by scraping, and genomic DNA was extracted. The relative abundance of each transposon mutant was estimated as described in Long et al. 2015.

### Microscopy and Damage Quantification

H&E-stained lung sections were imaged in bright field at 2X magnification on the Keyence BZ-X800. Images were processed identically within FIJI software (v2.3.0/1.53p) for image clarity. To quantify damage, an artificial neural network-based damage identification model was developed within QuPath (v0.3.2). Eight images (1 image per experimental group) were utilized for the single purpose of training the model, and the remaining 3 images per group were quantified using the model. To prevent bias, lung sections presented in this report were quantified as the closest to the mean damage of each experimental group.

### TnSeq Analysis

TnSeq libraries were prepared and transposon insertion counts were estimated as previously described (Smith et al. 2022). The H37Rv genome annotation used is NCBI Reference Sequence NC_018143.2. In total, 40 independent transposon libraries were sequenced after recovery from the 19 BXD genotypes and the two parent strains, B6 and D2. Two independent transposon libraries were placed under selection within each host genotype. However, in the cases of two BXD genotypes, BXD48a and BXD51, only one library could be sufficiently recovered for sequencing. Within TRANSIT (DeJesus et al. 2015), beta-geometric correction was used to normalize insertion mutant counts across all libraries, and pseudocounts were implemented. Insertion counts were totaled for each *Mtb* gene, and counts were averaged between individual replicate libraries per mouse. Per gene mean values were compared to the grand mean then log_2_ transformed.

To control for multiple hypotheses genome-wide, 10,000 permutation tests were performed to generate a resampling distribution used to derive the significance of transposon mutant selection between *in vitro* and *in vivo* conditions. An *Mtb* gene was considered “essential” within a host if mutants lacking that gene experienced a log_2_ fold change (LFC) value of -0.5 or lower post-selection in that host. To identify *Mtb* genes whose fitness was sufficiently variable to enable QTL mapping, the dynamic range in LFC across the parental and BXD strains for any given bacterial gene had to be at least 0.63. This value is the minimum LFC range of a mutant that was significantly selected in at least one host genotype after resampling (Q < 0.05). Mutants used for QTL mapping also needed to be sufficiently represented in the library, which was accomplished by requiring 4 or more TA transposon insertion sites to be detected for each bacterial gene. Further, each gene included needed to be significantly selected in at least one host genotype before resampling (p < 0.05).

### Genotyping and QTL Mapping

Previously published BXD genotypes containing 7,321 total markers were leveraged for QTL mapping (Wang et al. 2016). Markers that were not diagnostic of genetic differences between B6 and D2 were filtered out resulting in a total of 7,314 markers. Genotype data and phenotype data, including lung and spleen burden and TnSeq mutant fitness profiles, were imported into R statistical software (version 4.1.1) and formatted for QTL mapping within R/qtl2 (version 0.28) (Broman et al. 2019). The Leave One Chromosome Out (LOCO) approach was leveraged to estimate kinship between strains as a covariate for QTL mapping. Because all screened mice were male and infected in one batch, no other covariates were used for the initial mapping. Logarithm of odds (LOD) scores were calculated using the linear mixed model in R/qtl2 to establish phenotypic associations with marker loci across the host genome. Trait-wise significance thresholds for QTL were established by 10,000 permutation tests. The significance of gene class overrepresentation among the mapped QTL in comparison to class representation in the whole *Mtb* genome was established using Fisher’s Exact Test.

## Results

### B6 and D2 mouse strains experience distinct chronic disease profiles after aerosol infection

B6 and D2, the parent strains of the BXD panel, have divergent responses to *Mtb* infection (Chackerian and Behar 2003). Canonically considered *Mtb*-resistant, the T_H_1-skewed adaptive immune response initiated by B6 mice is capable of restraining *Mtb* growth after ∼21 days, resulting in a mean survival time of ∼230 days (Mitsos et al. 2000). However, D2 mice experience low IFN-γ producing T cell influx to the lung at 21 days, when adaptive immunity should begin, at which point B6 and D2 begin to diverge in bacterial growth restriction (**Figure 1A & B**) and lung damage (**Figure 1C**-**E**) (Marquis et al. 2008). D2 macrophages become foamy and dysfunctional, making it difficult to contain and control bacterial growth in the lung (Chackerian and Behar 2003). By 12 weeks post-aerosol infection, D2 lungs exhibit hyperinflammatory and fibrotic responses leading to overt damage (**Figure 1F**-**H**) and an ultimate mean survival time of ∼110 days (Medina and North 1998; Keller et al. 2006). In addition to genotype-specific differences between these BXD parent strains, we also observe a sex effect independent of genotype, demonstrated by higher susceptibility of males of both genotypes (**Figure S1A & B**) (Tsuyuguchi et al. 2001; Dibbern et al. 2017). Further, we find that in contrast to burden, all groups reduce lung damage by week 12 except for D2 males (**Figure S1C**-**G**).

**Figure 1:**
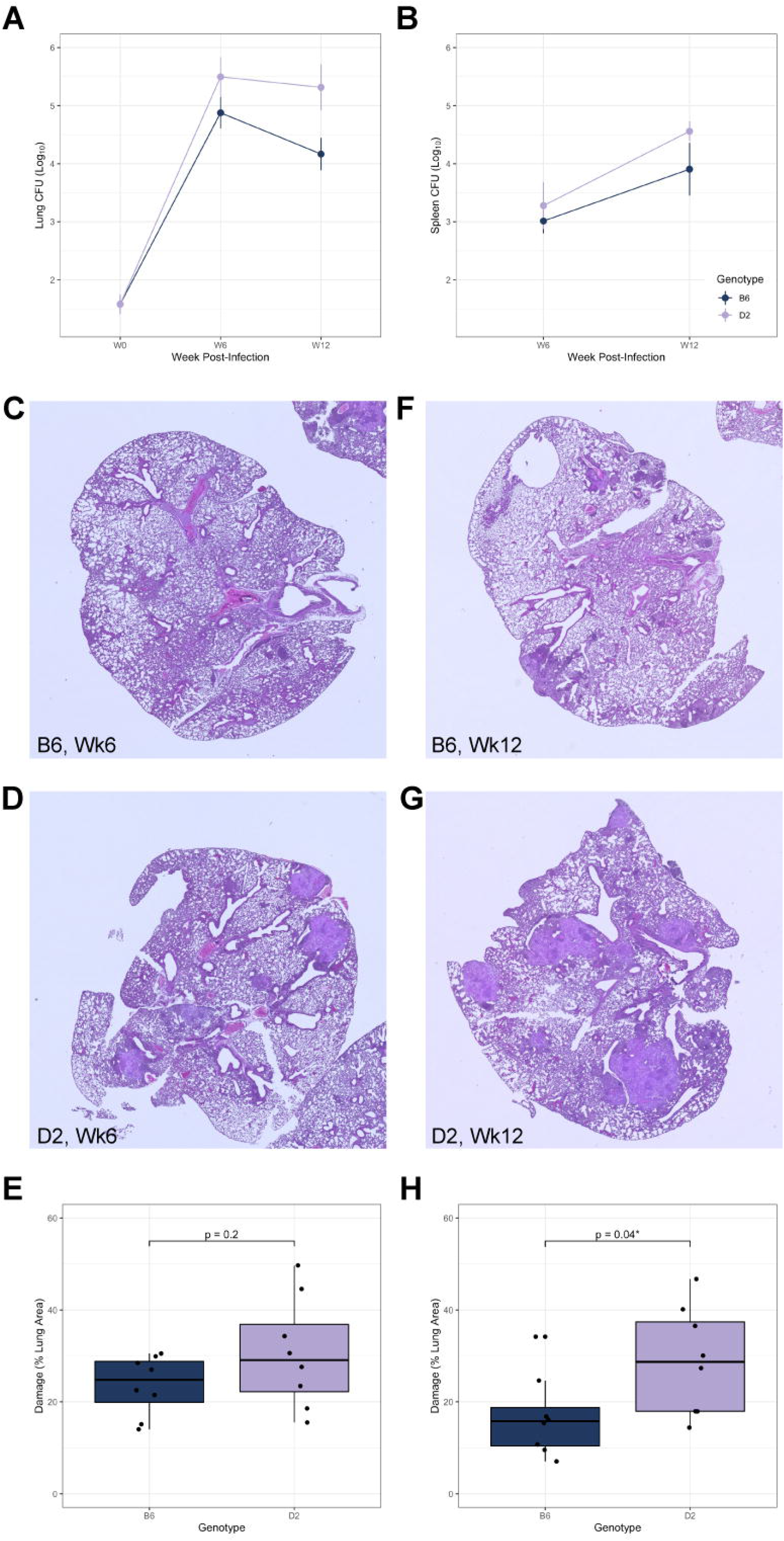
B6 and D2 are phenodeviant in TB susceptibility. (**A**) Log_10_ transformed colony forming units (CFU) recovered from lung tissue throughout the course of the aerosol infection (with an infectious dose of ∼40 CFU of *Mtb* H37Rv). (**B**) Log_10_ transformed CFU recovered from spleen throughout infection. (**C** & **D**) Male B6 and D2 H&E-stained lung sections taken 6 weeks post-infection, 2X magnification, representative of n = 4 per genotype. Corresponding female sections can be found in **Figure S1C** & **D**. (**E**) Lung damage was quantified by QuPath v0.3.2 using an artificial neural network-based damage identification model and reported as a percent of total lung section area (n = 4 per genotype & sex). Lung damage is not significantly different between B6 and D2 at 6 weeks post-infection by unpaired Student’s *t*-test. (**F** & **G**) Male B6 and D2 H&E-stained lung sections taken 12 weeks post-infection, 2X magnification, representative of n = 4 per genotype. Corresponding female sections can be found in **Figure S1E** & **F**. (**H**) Lung damage, quantified as in **E**, is significantly different between B6 and D2 at 12 weeks post-infection by unpaired Student’s *t*-test.

### Early clinical macrophenotypes fail to distinguish parental disease outcomes

With over 100 still extant recombinant inbred strains bred from B6 and D2 parental strains, the BXD family is a well-suited mammalian resource to map the phenotypic diversion during *Mtb* infection (**Figure 2A**). To study early disease traits and bacterial endophenotypes that predict differences in TB disease outcomes during chronic infection stages, we intravenously infected cohorts of the parental B6 and D2 lines and 19 BXD genotypes with a saturated TnSeq library of transposon (Tn) mutants. The Tn library contains knockouts of every *Mtb* gene that is not required *in vitro*, which collectively produces an infection similar to the wildtype infection strain (Sassetti and Rubin 2003; Smith et al. 2022). At 4 weeks post-infection, the parental B6 and D2 strains demonstrated approximately the same bacterial burden in lung and spleen (**Figure 2B**). Moreover, BXD strains did not exhibit sufficiently wide variation in bacterial burden at 4 weeks post-infection to precisely map the genetic cause of such a complex trait (**Figure 2C**; **Figure S2A** & **B**). Compound macrophenotypes, such as colony forming units (CFU) in organ homogenate, are highly polygenic, making it difficult to identify causal genes even with large panels of mice (Lavebratt et al. 1999; Mitsos et al. 2000; Mitsos et al. 2003; Yan et al. 2006).

**Figure 2:**
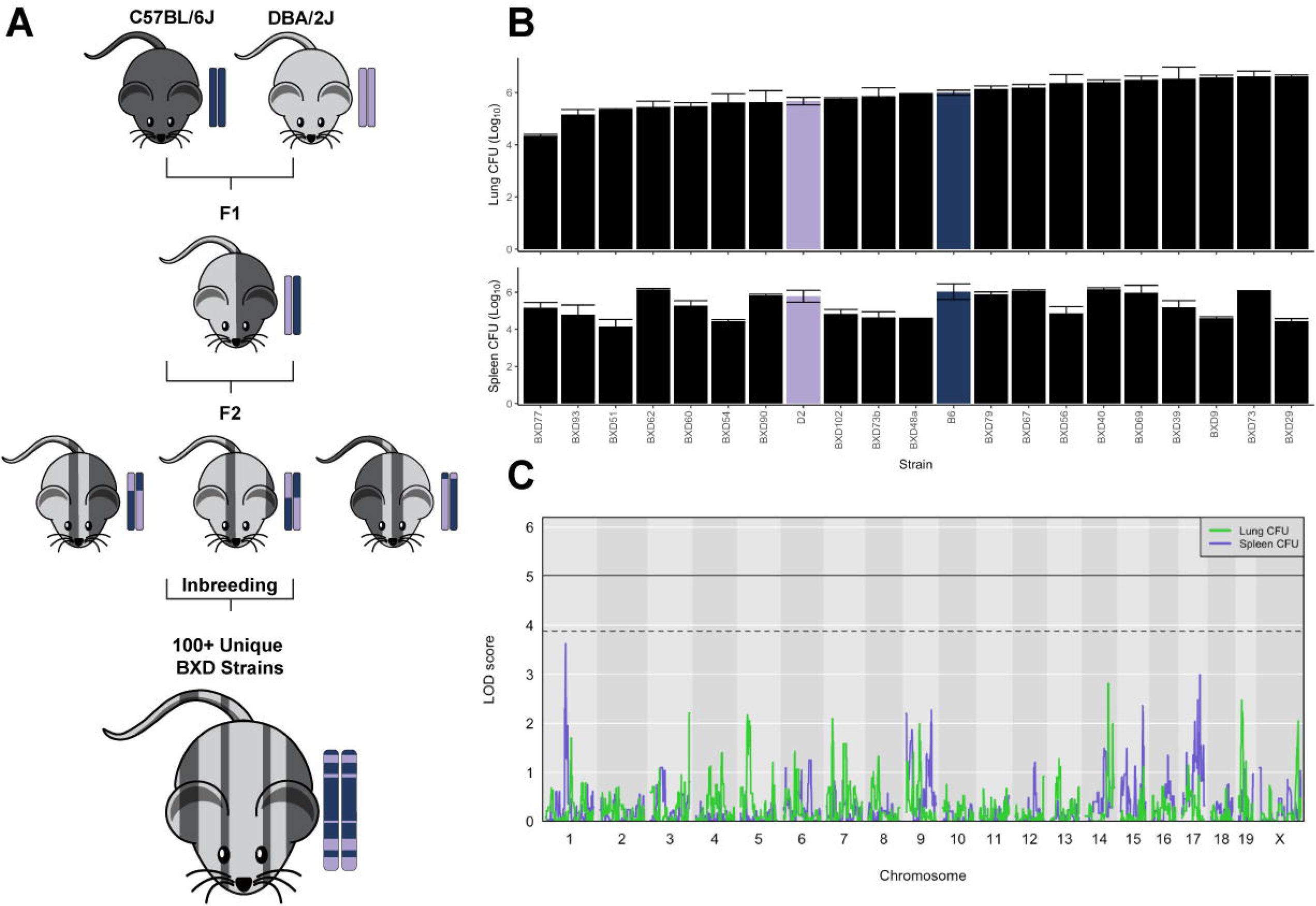
BXD panel exhibits natural genetic variation, but classical clinical traits cannot map genome-wide QTL at 4 weeks post-infection. (A) The BXD panel is a biparental recombinant inbred panel bred from C57BL/6J (B6) and DBA/2J (D2) and composed of over 100 mosaic strains. (B) Lung and spleen burden by host genotype (n = 1-5 per genotype). (C) QTL mapping of lung and spleen burden traits across the BXD cohort. Solid threshold represents a genome-wide significance of p = 0.05. Dashed threshold represents p = 0.20. Thresholds were calculated from 10,000 permutation tests.

### Transposon mutant fitness endophenotypes report on the host immunological microenvironment

To interrogate the biological mechanisms that predict immunodivergent outcomes, we quantified the abundance of each bacterial Tn mutant both before host infection with the TnSeq library and after recovery from mouse organs of each genotype. The log_2_ fold change (LFC) of abundance was calculated as a quantification of relative mutant fitness in each host condition. This LFC fitness value serves as a precise reporter of the host microenvironment and can be used to map host loci that impact *Mtb* mutant fitness (**Figure 3A**).

**Figure 3:**
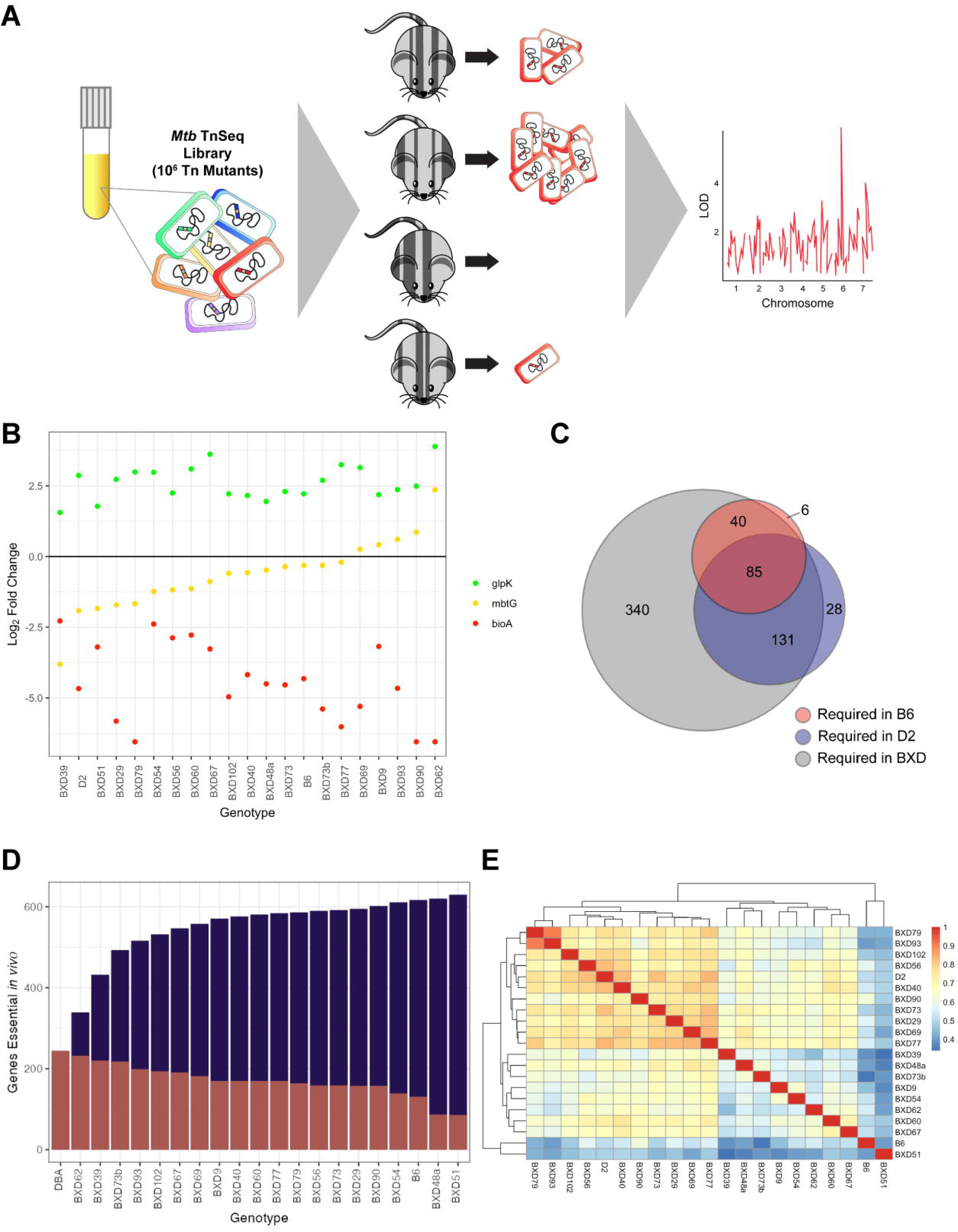
TnSeq mutant fitness is a sensitive reporter of the factors that drive disease outcome. (**A**) Multiple mice from each BXD and parental strain were infected with 10^6^ CFU of *Mtb* TnSeq library, representative of ∼10^5^ independent *Mtb* transposon mutants. For each mutant, the change between the initial abundance in the TnSeq library and the abundance recovered after 4 weeks of *in vivo* infection was compared to calculate a quantitative fitness metric (log_2_ fold change; LFC), which could be used downstream for QTL mapping. (**B**) Example plot showing an essential *Mtb* gene (*bioA*), a conditionally essential gene (*mbtG*), and a non-essential gene (*glpK*) across the BXD panel. (**C**) A Venn diagram depicting the overlap of essential genes between B6, D2, and the screened BXD strains. *Mtb* genes were deemed essential if the transposon mutant experienced a one-fold reduction (LFC < -0.5) within a given host genotype and were significantly different from *in vitro* conditions (p < 0.05) in at least one host genotype across the panel. (**D**) The number of *Mtb* genes essential (LFC < -0.5) for growth or survival in each diverse mouse strain across the panel (p < 0.05). Salmon indicates the mutants uniquely required for each host additional strain, and purple shows the cumulative requirement as each new host strain is added. (**E**) A heatmap depicting the total correlation of the library-wide bacterial fitness within each screened host strain.

Across the BXD panel, we identified three main classes of bacterial mutants, which are highlighted in **Figure 3B**. The first of these classes includes canonical *Mtb* genes that are essential for *in vivo* survival, exemplified by *bioA*. These genes remain essential for *Mtb* survival in both parental backgrounds and across the BXD strains (**Figure 3B**; “essential” profile in red). A second, “differentially required” set of mutants is exemplified in this figure by *mbtG*; the loss of *mbtG* did not significantly reduce bacterial fitness within B6 mice, but *mbtG* mutants experienced an extreme fitness reduction within D2 mice (**Figure 3B**; “differentially required” profile in yellow). Further, there is a spectrum of *mbtG* essentiality across the family, indicating that some variable host factor could modify the fitness of this mutant. In addition to *in vivo* essential and differentially essential *Mtb* genes, a third class of *Mtb* genes proved to be broadly dispensable for *Mtb* survival in these hosts, imparting an adaptive benefit for *Mtb* when the gene was disturbed, represented here by *glpK* (**Figure 3B**; “non-essential” profile in green).

When surveying the complete set of *Mtb* transposon mutants, we find that in comparison with B6 infection, *Mtb* requires almost twice as many genes to survive within D2 mice (**Figure 3C**). This finding suggests that there are detectable immunological pressures being exerted on *Mtb* within the highly inflammatory D2 line prior to the divergence in B6 and D2 bacterial control and disease outcome. This result replicates findings from previous work in which a greater number of *Mtb* genes were essential in T cell-deficient and IFN- γ-deficient mice than in B6 (Mishra et al. 2017; Smith et al. 2022). We additionally find that the BXD panel, while recapitulating most B6- and D2-essential genes, reveals additional subsets of genes that are only required within specific BXD strains (**Figure 3D**), highlighting the power of this genetically diverse panel to offer novel insights into the biology controlling TB outcome.

This spectrum of *Mtb* gene essentiality is further confirmed when we examine the correlation of the complete TnSeq library fitness between hosts (**Figure 3E**). Interestingly, the majority of the strains hierarchically clustered closer to D2 than to B6, indicating that they could be inheriting factors from D2 that put greater immunological pressure on *Mtb*.

### Transposon mutant fitness profiles map host-pathogen QTL (hpQTL)

We aimed to use these bacterial mutants as sensitive reporters of host immunological pressures within the early phase of infection at the onset of adaptive immunity. Using the LFC values collected for each Tn mutant population in the TnSeq library as quantitative endophenotypes (**Table S1**), we conducted QTL mapping. After removing non-dynamic genes for quality control (**Figure S3A**; *Mtb* genes that are not sufficiently varying in essentiality across the screened BXD genotypes), we found 140 genome-wide significant QTL, defined as p ≤ 0.05 after 10,000 permutations (**Table 1**). Of these QTL, 33 of the loci had a p-value ≤ 0.01 (**Figure 4A**).

**Figure 4:**
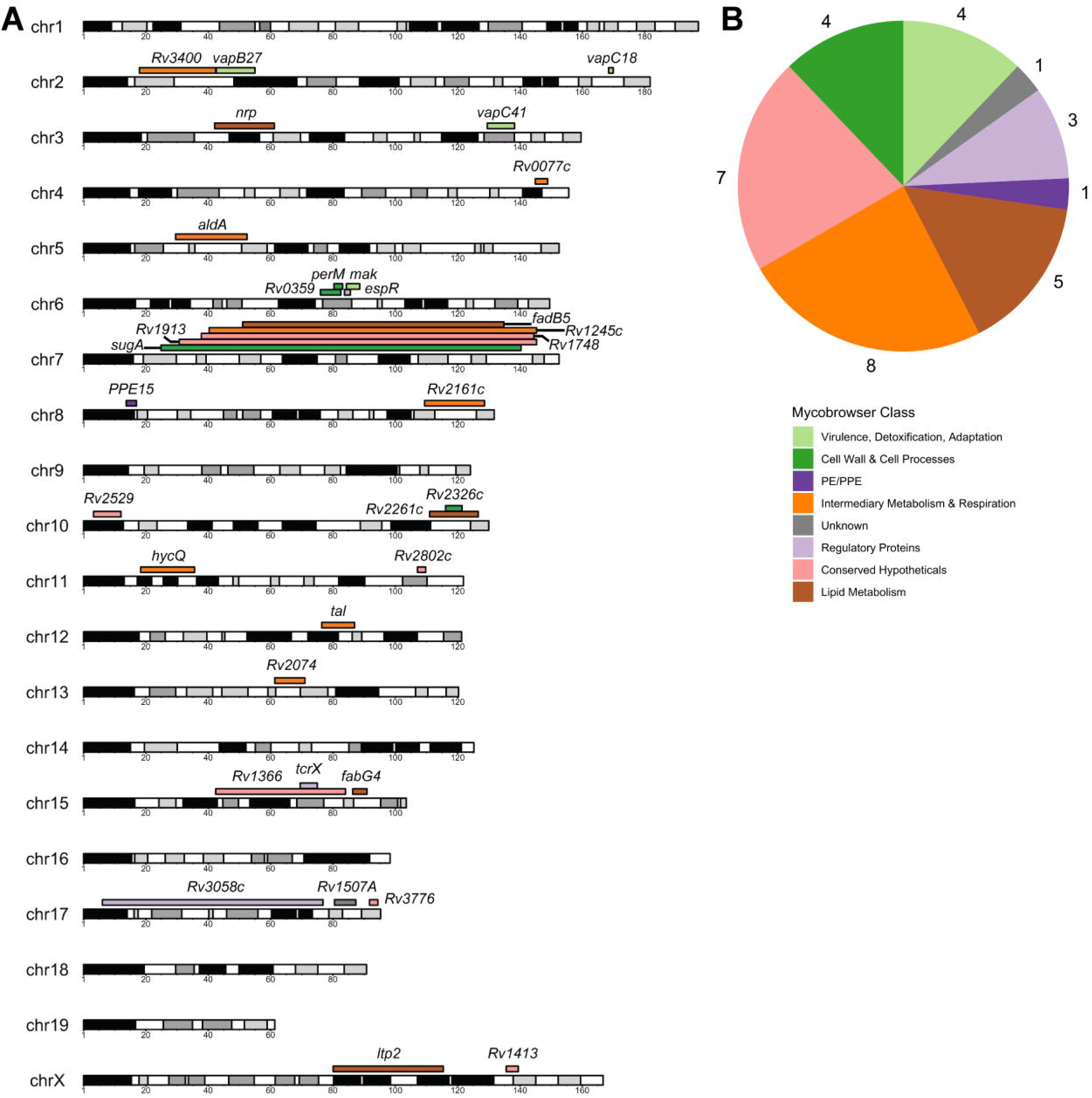
TnSeq mutant fitness endophenotypes map genome-wide significant QTL. (**A**) The QTL that reach a significance threshold of p ≤ 0.01. The color of the QTL corresponds to the Mycobrowser class of the gene absent from the transposon mutant that mapped the QTL. The width of each segment corresponds to the size of the 95% Bayesian confidence interval of the QTL. (**B**) A pie chart tabulating the Mycobrowser classes of the mutants that mapped highly significant QTL (p ≤ 0.01), beyond the genome-wide significance threshold of p ≤ 0.05.

**Table 1:**
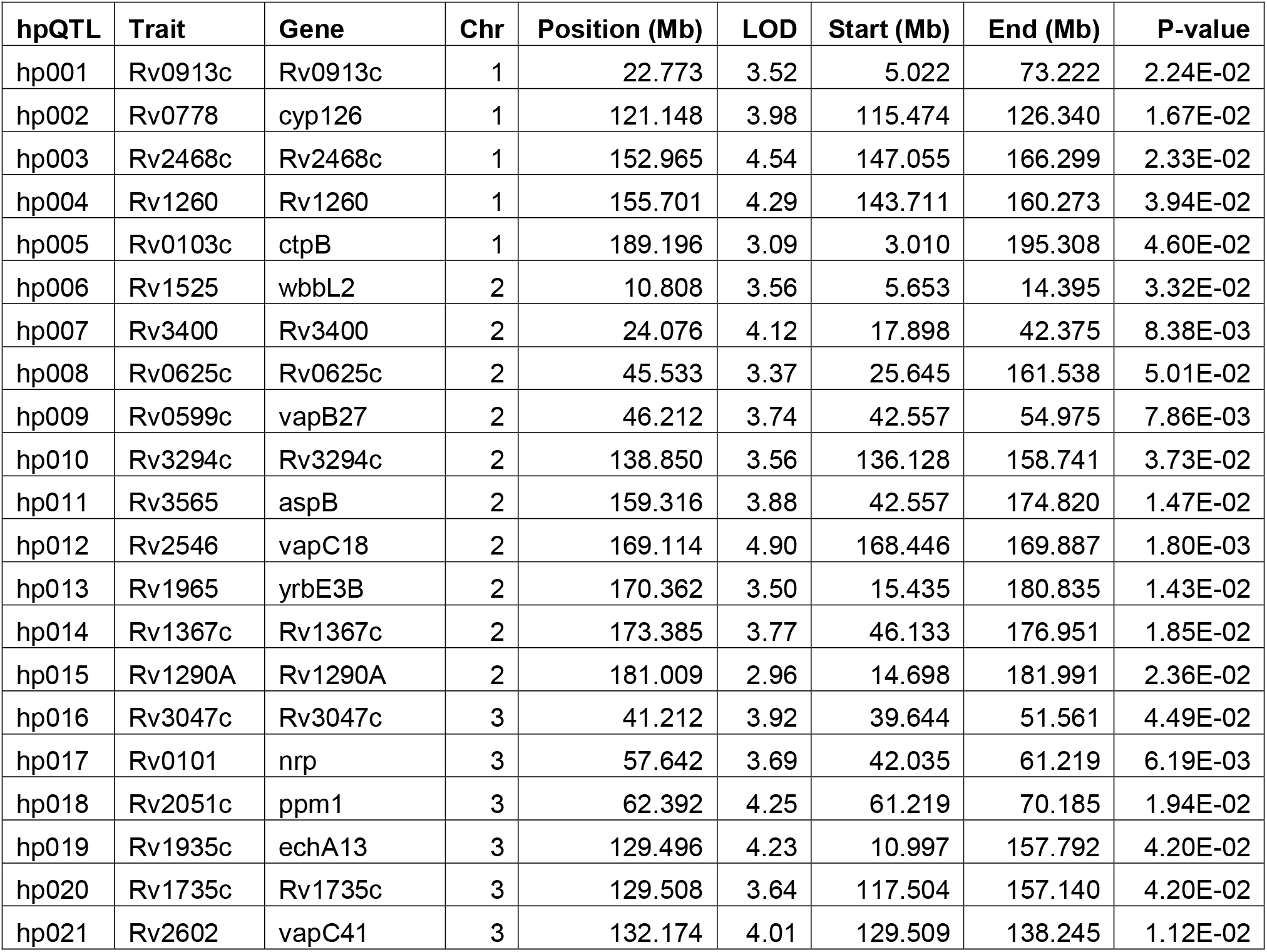
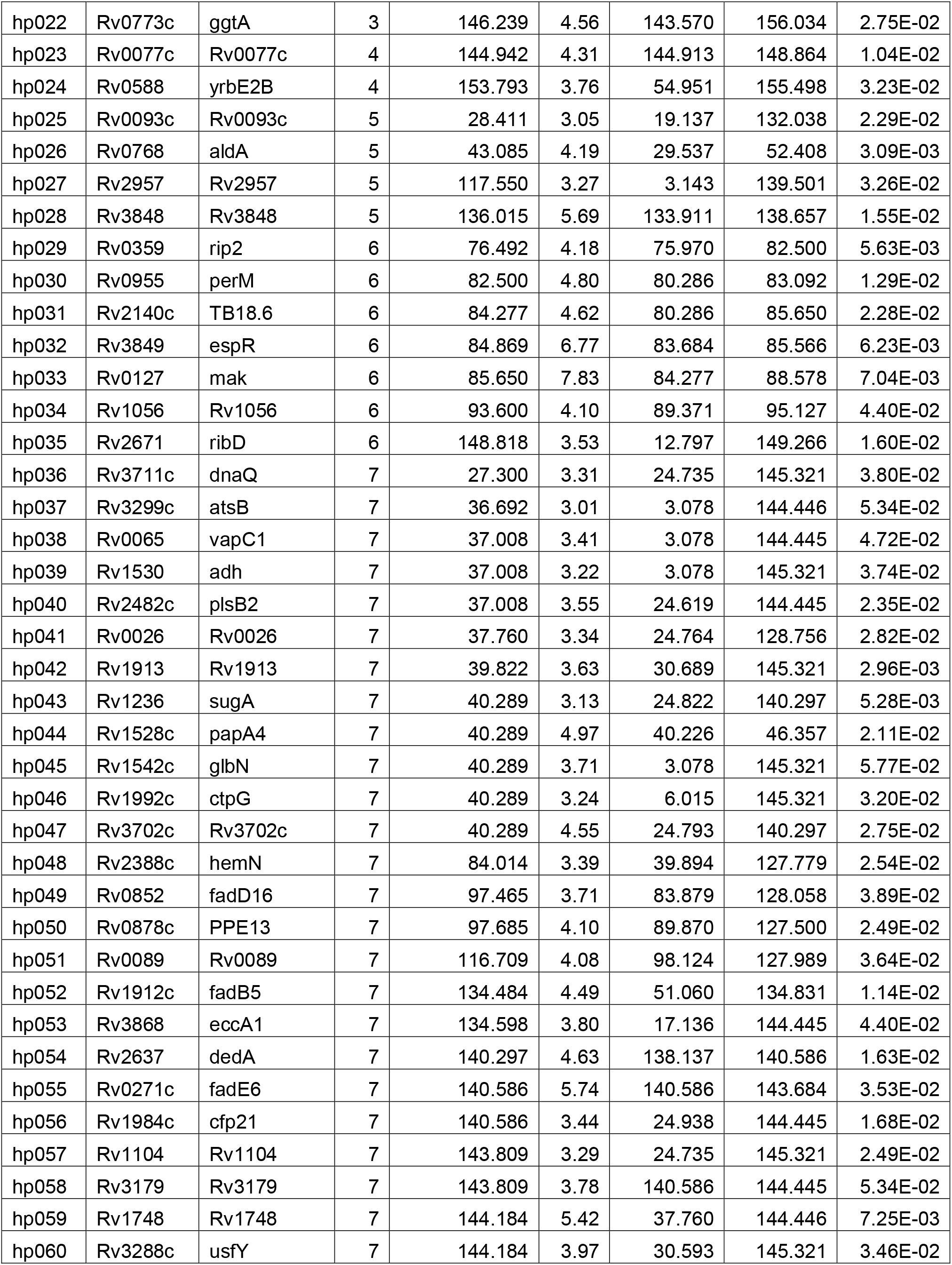
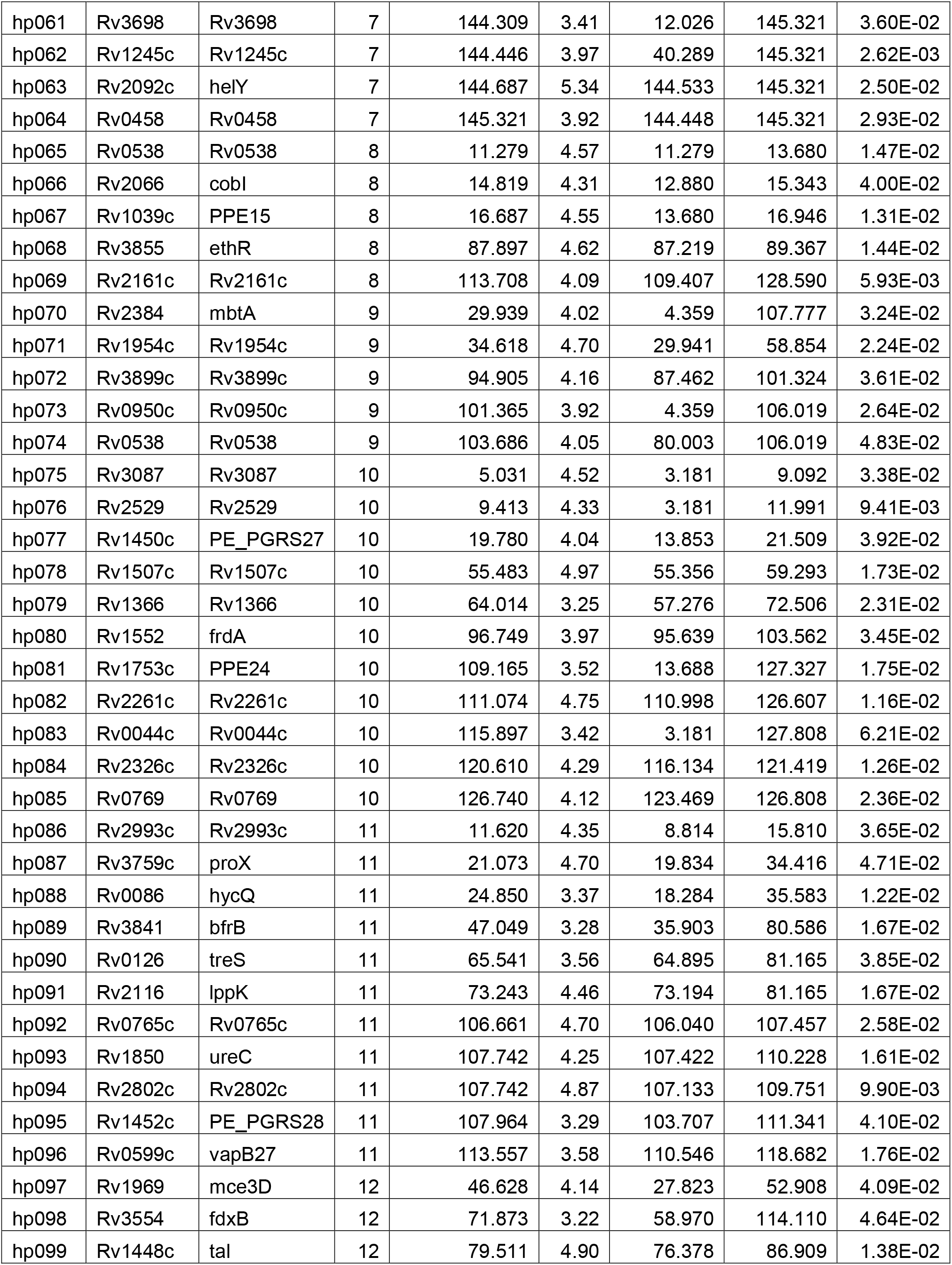
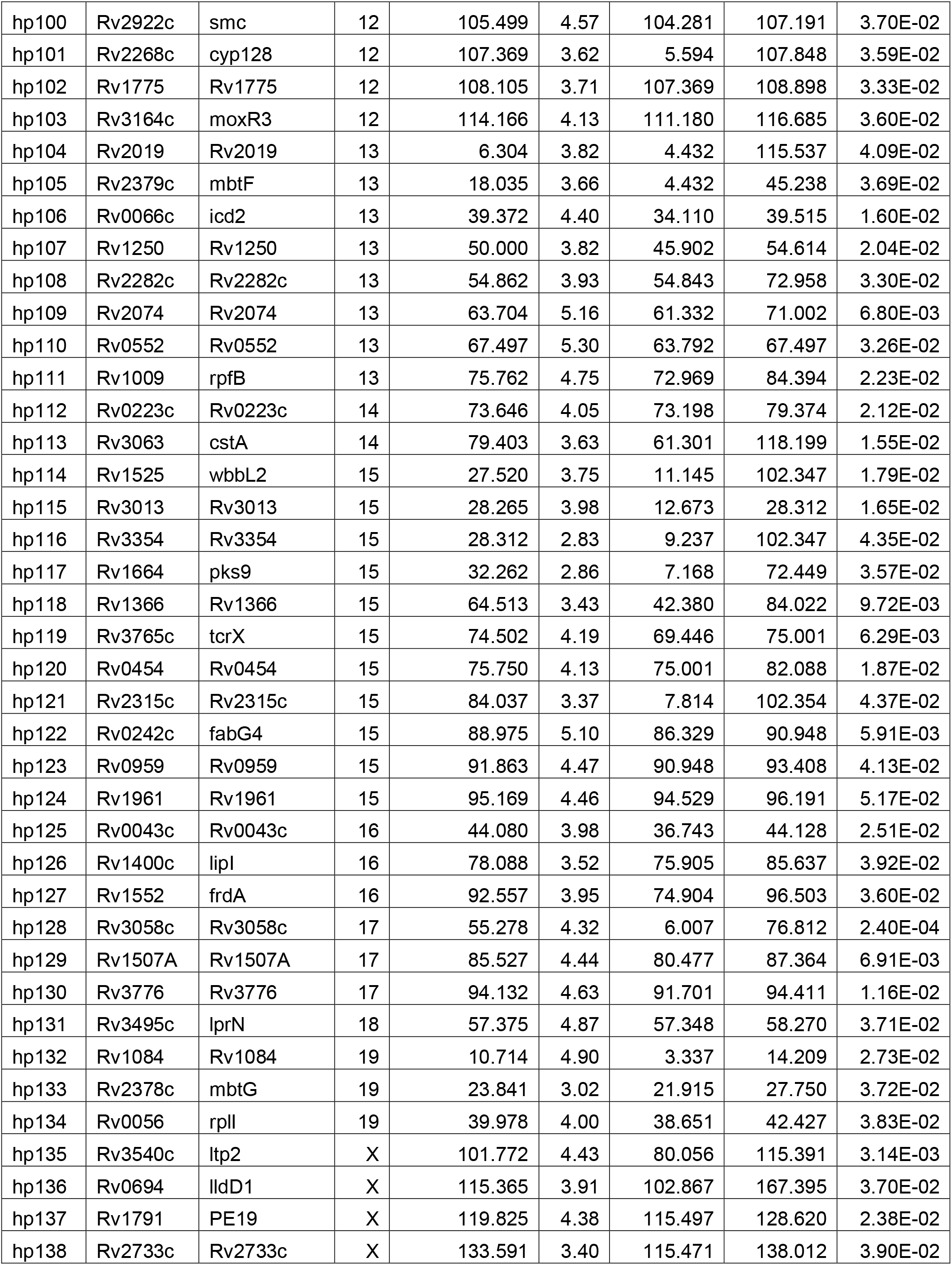
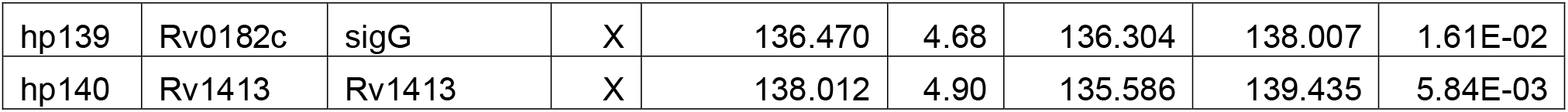
Genome-wide significant QTL (p≤0.05). “hpQTL” is the name for each host-pathogen QTL of genome-wide significance (p ≤ 0.05). “Trait” refers to the Rv number of the bacterial mutant whose fitness profile identified the hpQTL in the host genome. “Gene” is the bacterial gene annotation of the mutant if one is available on Mycobrowser. “Chr” is the host chromosome on which the QTL is located, and “Position (Mb)” is the position of the QTL in megabases. “LOD” is the maximum association score between the mutant fitness profile and the host locus. “Start (Mb)” and “End (Mb)” are the lower and upper boundaries of the Bayesian 95% confidence interval. “P-value” denotes the significance of the association between the bacterial mutant fitness and the host genetic locus.

Among the most significant QTL mapped in the screen, certain gene classes were marginally overrepresented in comparison to the genome-wide abundance of that class. Transposon mutants lacking genes that encode known virulence factors made up 12.1% of the top hits in this screen despite only representing 5.8% of the *Mtb* genome (p = 0.1) (**Figure 4B**) (Kapopoulou et al. 2011). Further, lipid metabolism genes made up 15.2% of the bacterial mutants mapping highly significant QTL while representing 6.6% of the *Mtb* genome (p = 0.07). Certain gene classes were abundant in approximately equal proportions in the mapped QTL as they were genome-wide, such as metabolism and respiration genes (22.8% among top QTL vs. 24.2% genome-wide prevalence) and genes encoding PPE/PE proteins (3.0% among top QTL vs. 4.1% genome-wide prevalence). Roughly half of *Mtb* protein coding genes remain unannotated (Whitaker et al. 2020), and 7 of the highly significant (p ≤ 0.01) bacterial fitness traits mapped by this screen correspond to bacterial genes with no known function. Taking these screen data in combination with previous *in vivo* TnSeq studies, these data will contribute to the de-orphanization of much of the *Mtb* genome.

### Multiple highly significant QTL map to host chromosome 6 “hotspot”

The nature of this screen enabled the discovery of host loci that impact the fitness of numerous bacterial mutants. Among the QTL mapped in this screen, 5 QTL mapped to chromosome 7 (**Figure 4A**), yet the 95% confidence intervals for each QTL span such a large part of the chromosome that identifying a causal host gene would be infeasible without further experimentation. Aside from the chromosome 7 locus, there appears to be a much narrower region on chromosome 6 (75.97–88.58 Mb) strongly linked to the essentiality of four bacterial genes: *Rv0127* (*mak;* sometimes annotated as *pep2*), *Rv0359* (*rip2*), *Rv0955* (*perM*), *Rv3849* (*espR*) (**Figure 5A-E**). For each of the mutants lacking these bacterial genes, apart from *perM*, the B6 sequence in this region is associated with lower bacterial mutant fitness (**Figure 5F-I**; **Figure S4A-E**). *Mtb* mutants lacking *perM* experience a fitness cost within genotypes that possess the *D* haplotype in this region.

**Figure 5:**
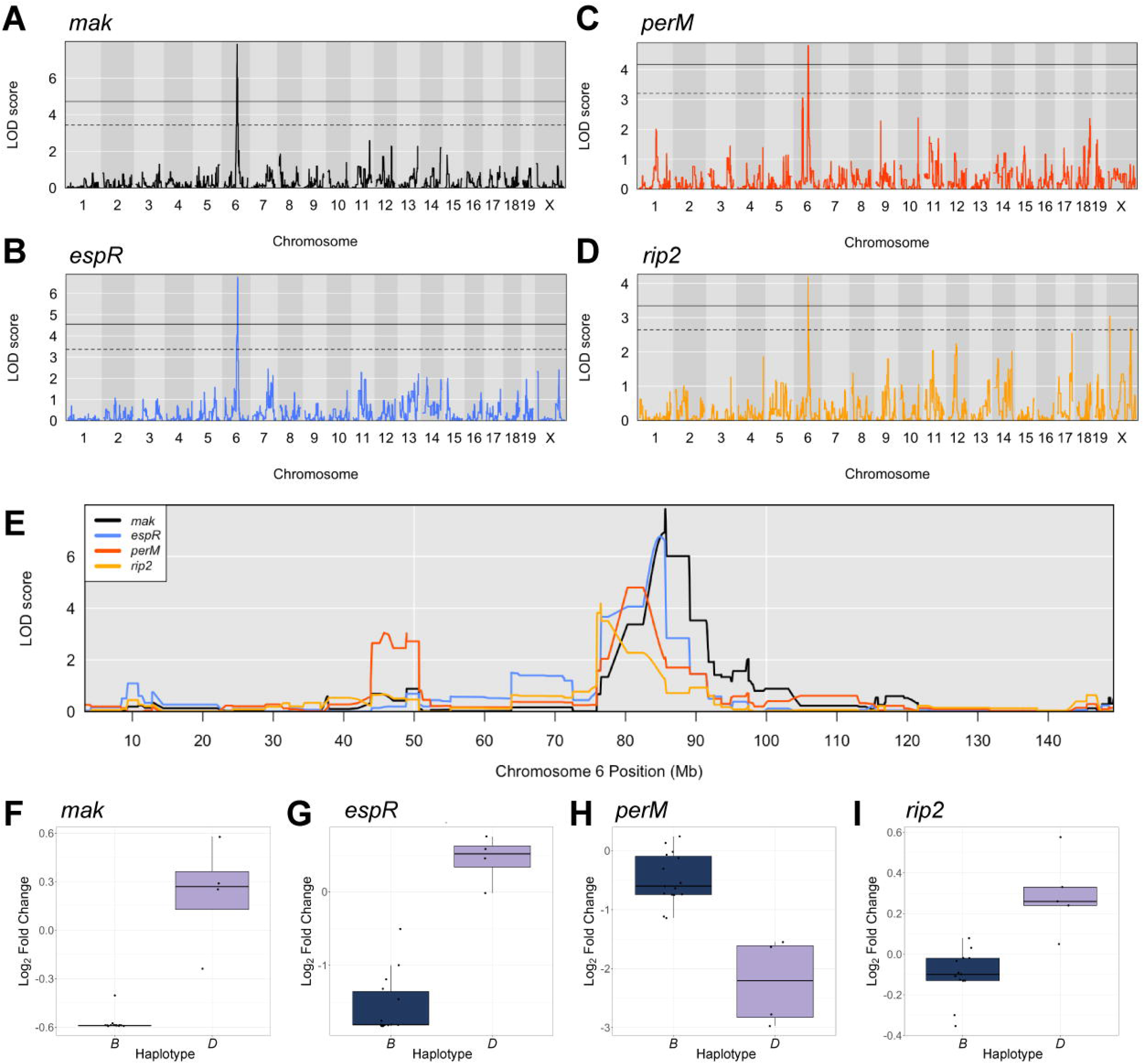
A chromosome 6 QTL hotspot highlights a genomic region controlling fitness of multiple bacterial mutants. QTL mapping of transposon mutants lacking (**A**) *mak,* (**B**) *espR*, (**C**) *perM*, and (**D**) *rip2*. After 10,000 permutation tests, the solid threshold represents p = 0.05, and the dashed threshold represents p = 0.20. (**E**) Chromosome 6 mapping overlap of the four transposon mutant fitness profiles that identified the QTL hotspot. (**F**-**I**) Boxplots representing the fitness of transposon mutants lacking *mak, espR*, *perM*, and *rip2* within BXD mice with *B* (navy) or *D* (lilac) haplotypes at the QTL position. BXD genotypes for which a haplotype state could not be called with 95% confidence were not included.

This QTL hotspot is mapped by *Mtb* genes that impact metabolism, virulence, and defense. Mak catalyzes the ATP-dependent conversion of maltose into maltose-1- phosphate as a part of the glycogen biosynthesis pathway that supports construction of the protective outer capsule of *Mtb* (Li et al. 2014). Of note, TreS, which converts trehalose to maltose upstream of Mak, also maps to a genome-wide significant QTL on chromosome 11 and has done so in previous *Mtb* infection screens (Smith et al. 2022). *rip2* is a putative zinc metalloprotease located in the *Mtb* cell membrane (Sklar et al. 2010), although very little is known about its function. PerM is an integral membrane protein that is known to play an essential role in enabling bacterial cell division under acid stress in B6 mice (Wang et al. 2019). Here, we report that *perM* mutants are even more severely attenuated within a D2 background, implying that PerM is differentially impacted by host immunological pressures. EspR is a key regulator of the ESX-1 secretion system (Rosenberg et al. 2011; Blasco et al. 2012), which itself plays a well-established role in mycobacterial virulence. Mutants lacking these bacterial genes map significant QTL within the interval of 75.97–88.58 Mb on mouse chromosome 6.

The 95% Bayesian confidence intervals of the QTL mapped by *rip2* and *perM* mutants overlap while the confidence intervals of *mak* and *espR* mutants overlap separately from the *rip2* and *perM* QTL. To test for the independence of these two groups of nearby QTL, we re-mapped each of the QTL within the chromosome 6 hotspot, iteratively incorporating the haplotype possessed by each BXD strain at each QTL as an additive covariate. Dependent loci will not map if the haplotype of one locus is factored into the mapping pipeline as a covariate. Conversely, the ability of a QTL to achieve significance independent of the haplotype state at another locus suggests that the two loci arise from distinct causal genetic factors within the host. These tests suggested two causal host loci exist within this highly significant QTL hotspot: one host factor underlying the *rip2* and *perM* mutant fitness QTL and one underlying the *mak* and *espR* QTL (**Figure 6A-D**).

**Figure 6:**
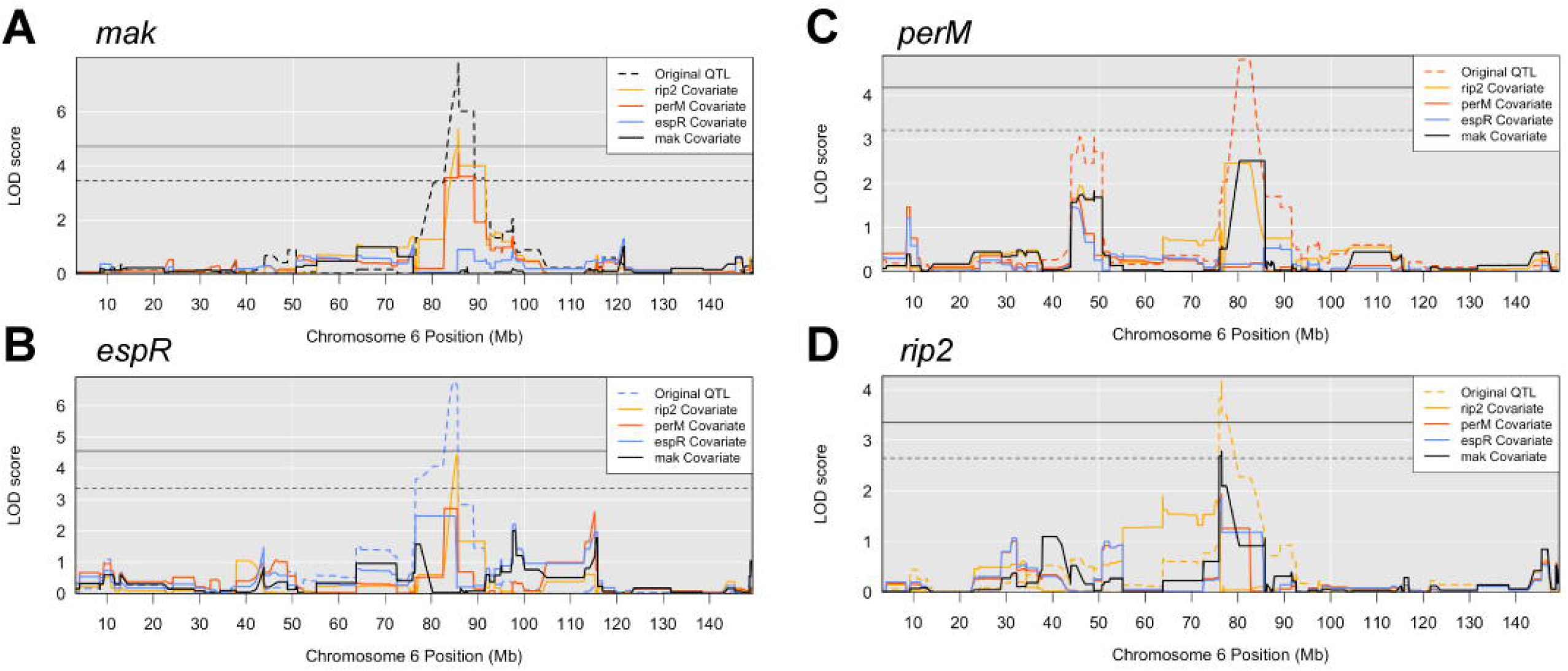
Independence tests identify two putative causal variants. Scan2 independence tests from R/qtl were used to identify whether the chromosome 6 hotspot may represent more than one causal locus. The QTL mapped by *mak* (**A**), *espR* (**B**), *perM* (**C**), and *rip2* (**D**) transposon mutants were remapped, incorporating the haplotype of the BXD mice at the locus of each other QTL as a covariate in the mapping analysis. The original QTL mapped by each mutant fitness profile are represented as dashed lines. The solid horizontal threshold represents p = 0.05 while the dashed horizontal threshold represents p = 0.20.

To identify host gene candidates that may mediate the survival of these two groups of bacterial mutants, we defined a pipeline to select for protein coding genes for which *D* haplotype has a unique SNP in comparison to *B* haplotype (**Figure 7A**). We passed each of the genes identified by this pipeline through a list of genetic and clinical criteria using independent but complementary data sets from a variety of *Mtb*-infected mammalian cohorts (**Figure 7B**) (Zak et al. 2016; Moreira-Teixeira et al. 2017; Ahmed et al. 2020). From this comparative analysis, we identified a putative host candidate within the region mapped by *mak*: MAX dimerization protein 1 (*Mxd1*), which is known to be upregulated in murine bone marrow-derived macrophages after infection with the hypervirulent *Mtb* HN878 strain (Roy et al. 2018). Further, we found a putative host candidate within the QTL mapped by *perM* mutants: ancient ubiquitous protein 1 (*Aup1*). Altogether, we have leveraged complementary screen data to identify plausible host gene candidates underlying these host-pathogen QTL.

**Figure 7:**
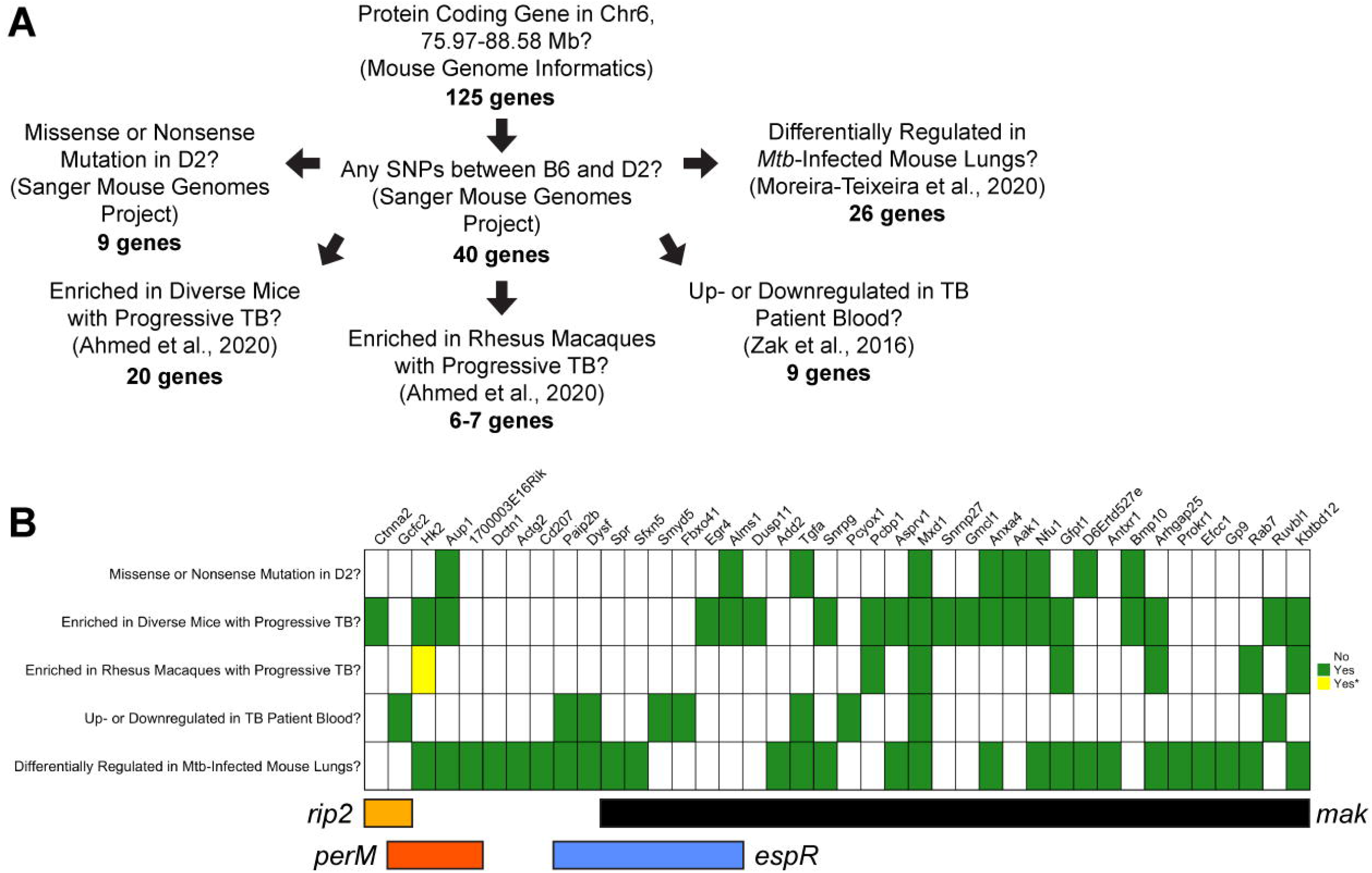
A bioinformatic pipeline highlights putative candidate genes underlying the QTL on chromosome 6. (**A**) A graphic summarizing the bioinformatic pipeline used to prioritize host gene candidates within each QTL. (**B**) A heatmap representative of the per-gene outcome of five distinct criteria: *i)* whether or not D2 possesses a missense or nonsense mutation in comparison to B6 according to the Wellcome Sanger Institute Mouse Genomes Project (Sanger MGP) (Adams et al. 2015), *ii)* whether the gene is significantly enriched in genetically diverse mice progressively infected with *Mtb* (Ahmed et al. 2020), *iii)* whether the gene is significantly enriched in Rhesus Macaques progressively infected with *Mtb* (Ahmed et al. 2020), *iv)* whether the gene is significantly up- or downregulated in TB patient blood (Zak et al. 2016), and *v)* whether the gene is differentially expressed in mouse lungs across variable host genotypes, *Mtb* genotypes, and doses (Moreira-Teixeira et al. 2020). If the answer is no, the space remains blank. Green indicates yes, and yellow indicates yes but cautions that the Rhesus Macaque gene is not a high confidence homolog. Only protein coding genes with a D2 SNP (as per Sanger MGP) that met at least one of the criteria are included in this visual.

## Discussion

While it is thought that nearly a quarter of the global population is infected with *Mtb* (Houben and Dodd 2016), only 5–10% of *Mtb* infections ultimately result in active TB (Gopal et al. 2014). This paradox has motivated decades of study on human genetic variants that could predict such a lethal prognosis in affected individuals. While GWAS among infected populations has identified numerous host loci associated with increased susceptibility to disease, a limited ability to dissect biological mechanism leaves many questions unanswered (Saul et al. 2019). QTL mapping, a parallel systems genetics technique, has been utilized for decades to identify host loci in genetically tractable populations linked to TB susceptibility; *Sst1* (susceptibility to tuberculosis 1), one of the most well-known TB susceptibility loci, was first discovered through a QTL mapping study in mice (Kramnik et al. 2000). Despite the widespread use of disease traits, including mean survival time, body weight, and bacterial burden, to identify host determinants of disease progression (Lavebratt et al. 1999; Kramnik et al. 2000; Mitsos et al. 2000; Mitsos et al. 2003; Yan et al. 2006), these phenotypes can be challenging to dissect immunological features that drive distinct *Mtb* pathogenesis in diverse hosts. Here, we describe a screen in which we leveraged the reproducibility, host variation, and phenotypic divergence of the BXD family and the molecular precision of the TnSeq library to identify novel axes of host-pathogen interaction during *Mtb* infection. We demonstrate the late (12 week) phenodivergence of the parental B6 and D2 mice, however we found that CFU serves as a weak predictor of host outcome at the onset of adaptive immunity 4 weeks post-infection. By utilizing TnSeq as a refined infection trait, we identified bacterial mutants that could distinguish the resistant B6 and susceptible D2 strains at the early 4 weeks infection timepoint. Finally, we leveraged the differentially required mutants as traits to conduct QTL mapping and identified 140 genome-wide *hp*QTL and a QTL hotspot encompassing a cluster of *Mtb* virulence and cell wall genes. Using this BXD TnSeq platform and a novel candidate prioritization pipeline, we identified viable host gene candidates for future study.

This screen highlights the importance of intentionality in design for deep phenotyping platforms to optimally identify the causal factors underlying gene associations. In contrast to complex macrophenotypes, which require hundreds of mice to sufficiently power causal locus identification for highly polygenic traits, molecular endophenotypes, such as *Mtb* transposon mutant fitness, allow us to learn more about the intricate details of *Mtb* invasion in the early stages of infection, which is particularly helpful in an experimental framework with fewer host genotypes. The strength of TnSeq as a deep phenotyping platform lies in the redundancy of each *Mtb* gene knockout; within the TnSeq library, each bacterial gene that is non-essential *in vitro* was perturbed at 4 or more unique transposon insertion sites, resulting in multiple subpopulations of unique replicates of each *Mtb* gene knockout. These mutant replicates taken together paint a robust picture of individual *Mtb* gene requirement *in vivo*, and multiple host replicates strengthen the reproducibility of each *Mtb* mutant fitness profile. In contrast with our previous TnSeq study that utilized nearly 70 host genotypes (Smith et al. 2022), this TnSeq study includes 21 unique genotypes and 73 mice, including the parental strains B6 and D2. The greater number of transposon mutant *hp*QTL identified in this screen demonstrates the power of a two-state model to segregate allele effects, as opposed to the eight-state model present in the Collaborative Cross recombinant inbred panel. We, however, cannot eliminate the possibility that the smaller number of host genotypes included in this study did not sufficiently limit statistical noise during QTL mapping analyses.

Using bacterial mutant profiles as endophenotypes, we identified 140 genome-wide significant (p ≤ 0.05) associations between *Mtb* transposon mutant survival and host genetic loci in a diverse cohort, which may contribute to divergence in clinical outcomes between *Mtb*-resistant and susceptible hosts. These host-pathogen QTL (*hp*QTL) and the transposon mutant fitness profiles across the BXD panel have been shared with GeneNetwork.org (Chesler et al. 2004; Parker et al. 2017; Ashbrook et al. 2021), an online compendium of BXD phenotypes. Increasing the precision of bacterial traits creates a rich environment for screen-based hypothesis generation in a smaller host population, thereby lowering the cost through lower animal count and less total infection time. This screen serves as a proof-of-concept that large host populations are not an absolute necessity to uncover novel and biologically meaningful axes of the host-pathogen interface. Molecular endophenotyping enables a much more accessible experimental platform to assist in our ongoing struggle against *Mtb*.

From this discovery screen, we have identified several plausible host gene candidates. Variants in *Aup1* represent a possible axis for interaction between *Mtb* and the mammalian host. The acetyltransferase activity of *Aup1* has been shown to be exploited by flaviviruses to trigger autophagy of lipid droplets within the cell, which produces ATP for the virus, thereby promoting viral replication (Zhang et al. 2018). Although not much is known about whether *Mtb* interacts directly with this protein, *Mtb* is one of the few bacteria capable of generating lipid droplets (Daniel et al. 2011). Host lipid droplets, specifically those derived from foamy macrophages, have been shown to promote the production of defensive cytokines during *Mtb* infection (Jaisinghani et al. 2018), implicating a potentially intriguing locus for host-pathogen interaction. The other high priority candidate *Mxd1* encodes the protein MAD, which competes with MYC to bind with MAX, forming a transcriptional repressor. This mechanism has led *Mxd1* to be considered a putative tumor suppressor gene. *Mxd1* has been shown to be upregulated in murine bone marrow-derived macrophages upon initial infection with the hypervirulent *Mtb* HN878 strain (Roy et al. 2018) and to regulate the fitness of murine dendritic cells (Anderson et al. 2020), yet the function of *Mxd1* in response to *Mtb* has yet to be determined. Together these candidates provide feasible next steps for mechanistic interrogation of these loci.

While the functions of many mycobacterial genes remain unknown (Whitaker et al. 2020), this transposon mutant screen within the BXD family builds directly upon a previous screen in the Collaborative Cross panel (Smith et al. 2022) to further articulate the conditional necessity of many canonically “non-essential” *Mtb* genes within a spectrum of host microenvironments. A substantial number of *Mtb* genes known to be non-essential in B6 mice prove to be essential in a subset of these genetically diverse hosts, suggesting that these bacterial genes play an active yet undiscovered role in pathogenesis under specific immunological conditions. These new host conditions have been shared with MtbTnDB (Jinich et al. 2021), a central repository encompassing over 60 published *Mtb* TnSeq screens for ease of browsing and inter-screen comparison. For mycobacterial researchers exploring *Mtb* genes traditionally thought to be non-essential *in vivo*, this work challenges such a notion and offers a new model that will provide insight into the host-pathogen dynamic underlying this conditional *in vivo* essentiality.

## Data Availability Statement

All relevant data to support the findings of this study are located within the paper and supplementary files. Genome sequence data is being deposited in the NCBI Gene Expression Omnibus (GEO). Transposon mutant fitness profiles within each BXD genotype and parental genotypes will be made publicly available on MtbTnDB (https://www.mtbtndb.app/). BXD phenotypes are available on GeneNetwork (https://www.genenetwork.org).

## Acknowledgements

We thank Anna Tyler, Matthew Mahoney, and Greg Carter for their assistance in concept development; Emily Hunt, Kris Riebe, and Summer Harris for technical assistance; Erin Curtis for graphic design and thoughtful discussions; Tobin and Smith Lab members for constructive feedback; Arthur Centeno for BXD phenotype import to WebQTL; and Douglas Marchuk for insightful comments on the manuscript. We are especially grateful to Richard Baker for rigorous biostatistical suggestions, meticulous data management and analysis, and conceptual development. Biocontainment work was partially performed in the Duke Regional Biocontainment Laboratory, which received partial support for construction from the National Institutes of Health, National Institute of Allergy and Infectious Diseases (UC6-AI058607; G20-AI167200).

