## Supplemental Figures S1-S4 for "Genome-wide screen identifies host loci that modulate *M. tuberculosis* fitness in immunodivergent mice"

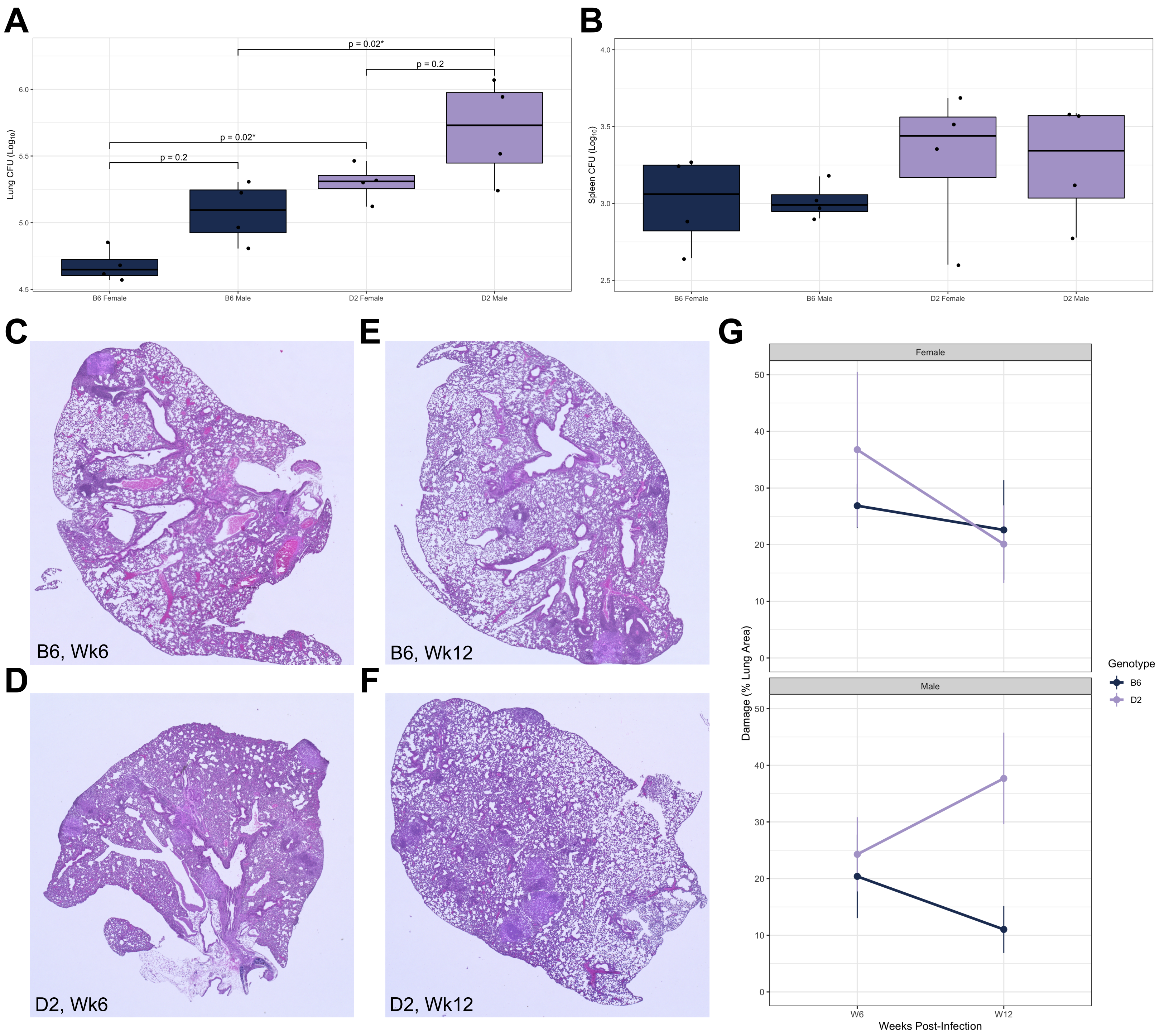


**Figure S1: Sex has a significant effect on susceptibility by aerosol but is not dependent upon host genotype.** (**A**) Both sex (p = 0.007) and genotype (p = 0.0003) have a significant effect on lung burden at 6 weeks post-infection by ANOVA. p-values included in the plot were calculated by Tukey's *posthoc* test. (**B**) Neither sex nor genotype significantly impacted spleen burden at 6 weeks post-infection by ANOVA. (**C-F**) Female B6 and D2 H&E-stained lung sections taken 6- and 12-weeks post-infection, 2X magnification, representative of n = 4 per genotype per timepoint. (**G**) Damage quantification of H&E-stained lung sections in QuPath v0.3.2 using an artificial neural network-based damage identification algorithm (n = 4 per genotype, sex, and timepoint).

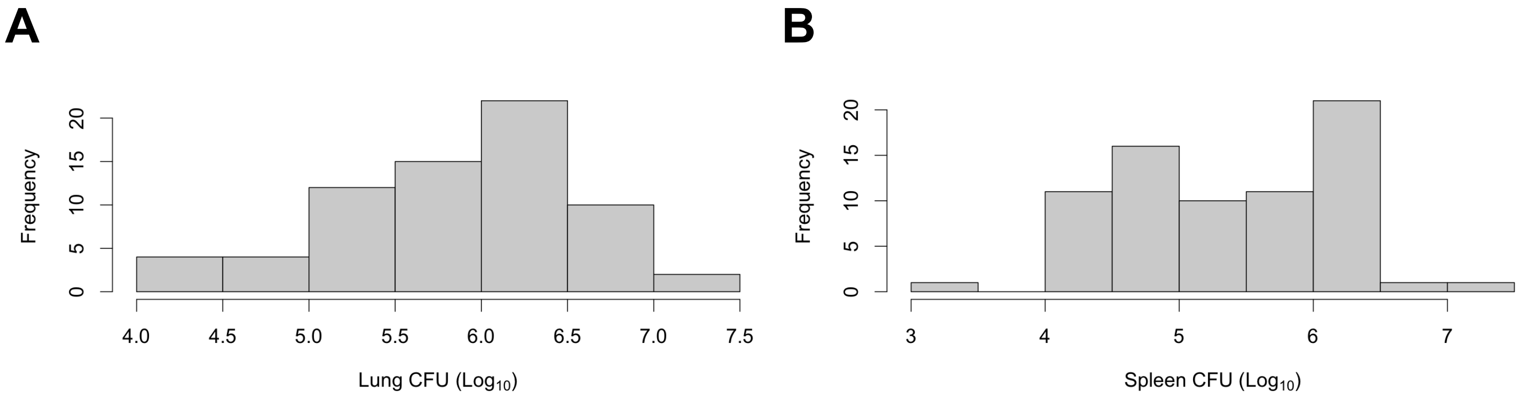


**Figure S2: Distribution of lung and spleen burden across the BXD panel at 4 weeks post-infection.** Histograms of lung (**A**) and spleen (**B**) burden values quantified across each screened BXD individual.

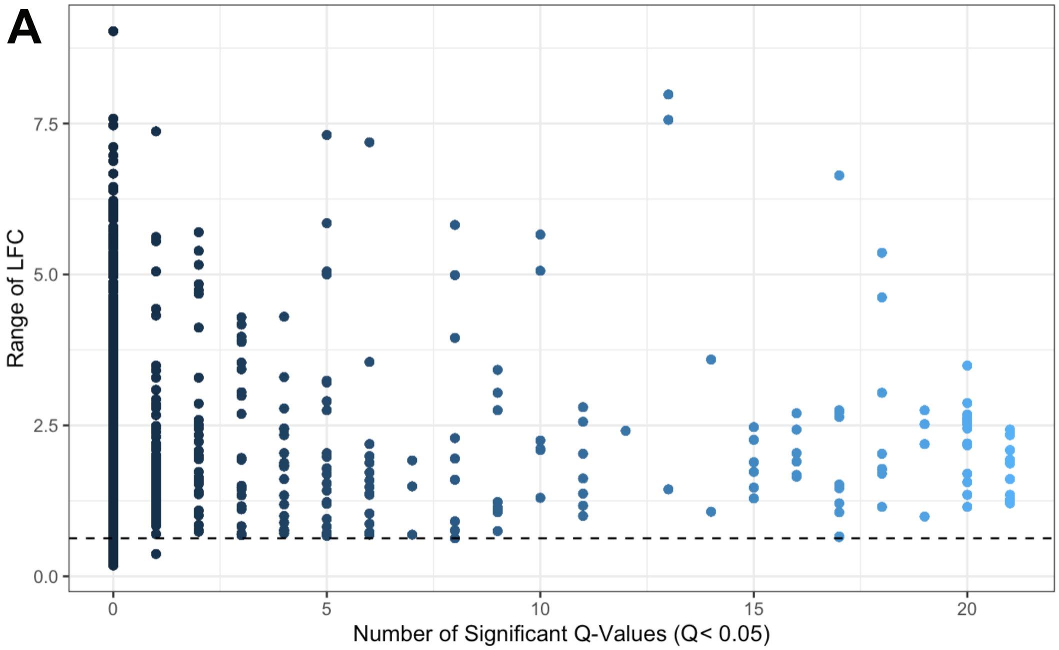


**Figure S3: Thresholding criteria of transposon mutant fitness profiles for QTL mapping.** To conduct QTL mapping on sufficiently varying bacterial genes, we excluded transposon mutants with a dynamic range of less than or equal to 0.63 $\log_2$ fold change (LFC), which is the minimum range of *Mtb* mutants with at least one significant Q-value between *in vitro* and *in vivo* conditions across the panel.

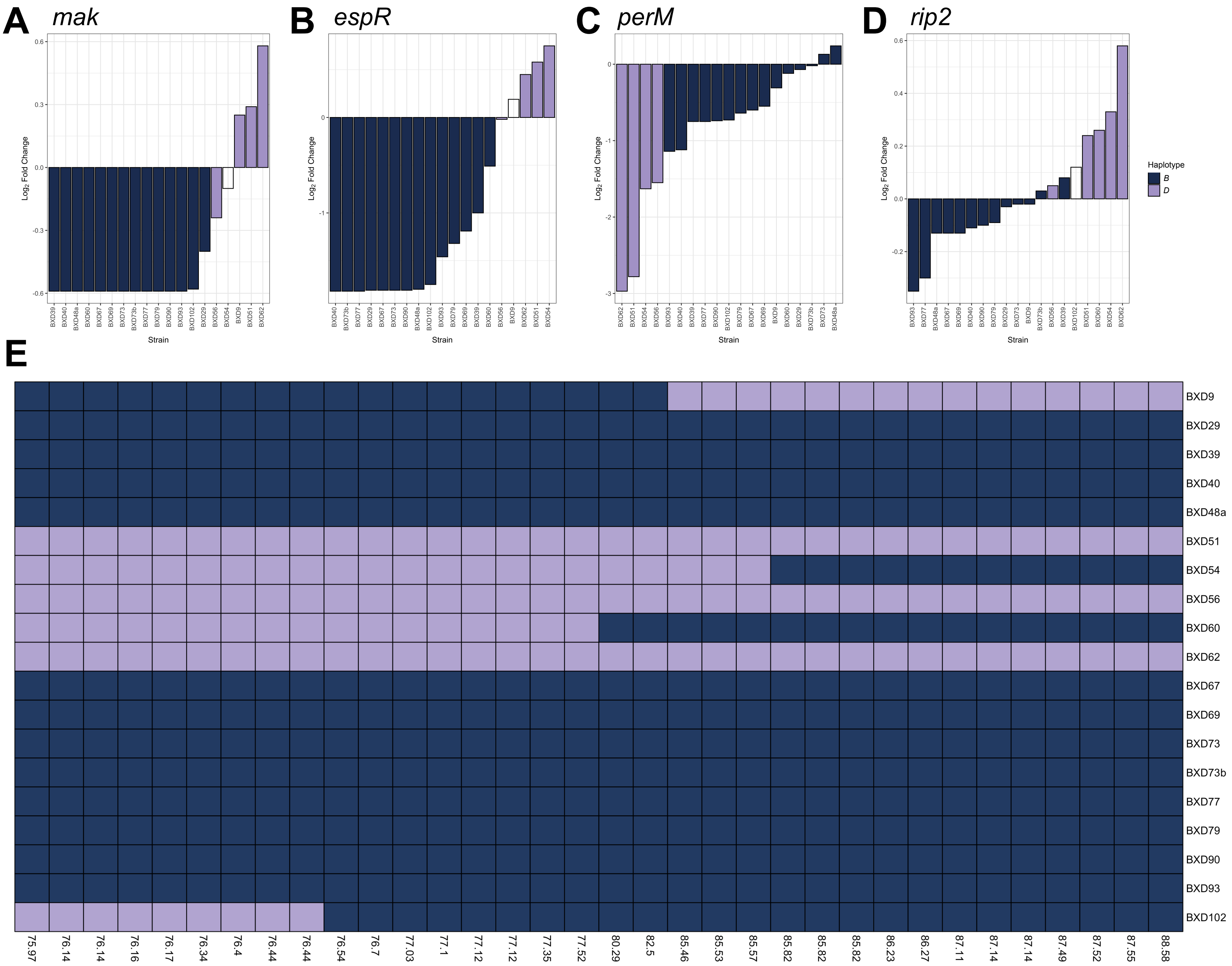


**Figure S4: BXD haplotypes within the chromosome 6 QTL hotspot.** Bar plots comparing the predicted haplotypes of each BXD genotype at the QTL position for (**A**) *mak*, (**B**) *espR*, (**C**) *perM*, and (**D**) *rip2* with the fitness of each transposon mutant within each BXD genotype. Empty bars represent haplotype states that could not be assessed with at least 95% confidence at the QTL. (**E**) A visualization of the BXD genotypes across the hotspot interval. Position values are represented in Mb.
